# A one-tube method for rapid and reliable plant genomic DNA isolation for PCR analysis

**DOI:** 10.1101/2020.02.13.948455

**Authors:** Wei Hu, J. Clark Lagarias

**Affiliations:** Department of Molecular and Cellular Biology, University of California, Davis, CA 95616 USA

## Abstract

**Background:** Consistent isolation of high quality plant genomic DNA is a prerequisite for successful PCR analysis. Time consumption, ease of operation and procedure cost are important secondary considerations for selecting an effective DNA extraction method. The simple, reliable and rapid DNA extraction method developed by Edwards and colleagues in 1991 [1] has proven to be the gold standard.

**Results:** Through modification of the Edwards method of extraction, we have developed a one-tube protocol that greatly improves the efficiency of plant DNA extraction and reduces the potential for sample contamination while simultaneously yielding high quality DNA suitable for PCR analysis. We further show that DNA extracts prepared with this method are stable at room temperature for at least three months.

**Conclusion:** The one-tube extraction method yields high quality plant DNA with improved efficiency while greatly minimizing the potential for cross contamination. This low-cost and environment-friendly method is widely applicable for plant molecular biology research.

## Background

Preparation of plant genomic DNA for PCR analysis is routinely performed in plant biology research laboratories. Among the many DNA extraction protocols in the literature, the method developed by Edwards *et al* [1] is the most widely used, as evidenced by its near three thousand citations. The popularity of this method relies on its simplicity, the generation of consistently high quality DNA, and its reproducible PCR amplification. Although alternative and more rapid protocols have been developed, the quality of DNA is often sacrificed for the reduction in preparation time using these protocols [2, 3]. Failed PCR analyses are not only disappointing, but often require the researcher to repeat the entire experiment. Thus, the reliability of the Edwards method has consistently outweighed the apparent time-saving benefits of other more rapid procedures.

## Results and Discussion

To improve the efficiency of the already simple and rapid Edwards method for genomic DNA extraction for PCR analysis [1], we added isopropanol directly with the crude tissue homogenate to precipitate DNA in the original extraction vessel (see methods). Since genomic DNA can be recovered from pelleted cellular debris, this modification obviates the need for a new tube for isopropanol precipitation of DNA. In this regard, the supernatant transfer and labeling of a secondary tube in the Edwards method is a time-consuming step that has the potential for mislabeling DNA samples when many samples are being prepared simultaneously. This modification is thus environmentally friendly and avoids potential cross contamination, since a single tube is used throughout the extraction procedure. After the initial centrifugation, a rapid air-dry method was found to be preferable to vacuum drying to remove traces of extraction buffer and isopropanol. The precipitated genomic DNA was then resolubilized by adding water to the ground plant debris. The improved one-tube extraction protocol takes ten minutes to process two samples and is a considerable simplification of the experimental manipulations.

As shown in Figure 1, DNA extracted from leaves of *Arabidopsis thaliana* L. using the one-tube protocol was amplified with PCR efficiency similar to that extracted using the Edwards method for < 2 kb amplicons. Our extraction method was slightly less efficient for > 2 kb amplicons in routine PCR. These results indicate that the debris does not appreciably inhibit PCR amplification of the extracted DNA. We also compared our method with a one-step protocol developed by Kasajima *et al* [3], which is the most rapid DNA extraction method available. DNA extracted by this method was less efficiently amplified by PCR for amplicons smaller than 2 kb, and proved unable to support amplification of larger amplicons. Our one-tube method proved superior to that of Kasajima *et al*.

**Figure 1.**
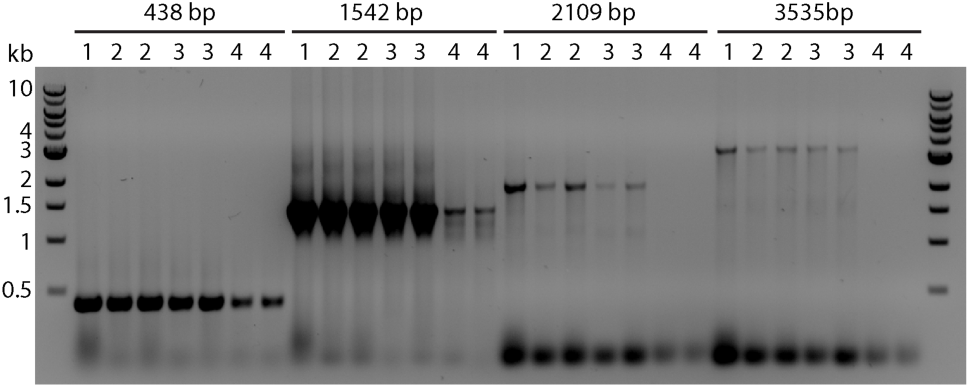
PCR analysis of DNA extracts prepared using the standard Edwards protocol (1), the one-tube protocol (2 and 3) and the one-step protocol of Kasajima *et al* (4). Two independent DNA extracts from (2) to (4) are shown; DNA extracts of (3) were stored at room temperature for three months.

To test the stability of DNA isolated by the one-tube method, DNA solutions were kept on the bench at room temperature and analyzed by PCR over time. Figure 1 shows that DNA extracts could be stored for at least three months at room temperature without a measurable reduction in PCR efficiency compared to freshly prepared DNA extracts. Refrigeration or freezing is thus unnecessary for storage of DNA extracted by this method. This is particularly convenient when multiple and sequential PCR analyses of the same DNA sample become necessary. For ground plant tissue that contains large amounts of fibrous material, e.g. mature siliques, precipitated debris may float up during prolonged storage. In this case, a brief centrifugation will re-establish the debris and DNA supernatant phases with no loss of PCR amplification efficiency.

We further validated that DNA extracted by the one-tube method permits amplification of large DNA targets using long-range PCR technology with an efficiency similar to DNA prepared with the Edwards method (Figure 2). In particular, a targeted ∼10 kb fragment was steadily amplified, indicating that DNA of high molecular weight was preserved throughout the extraction procedure. Figure 3 shows that reliable PCR amplifications of selected targets were achieved with DNA extracted from different *Arabidopsis* tissues using the one-tube method. In addition, this method worked well for a wide range of leaf biomass input from 1 to 100 mg (representing a single cotyledon or two rosette leaves) with similar efficiency of PCR amplification. This indicates that the extraction volume of 200 μl used in the protocol is sufficient regardless of the amount of tissue up to 100 mg. This is particularly helpful for extraction of pooled tissue samples from multiple plants. Grinding more than 50 mg of leaf in 200 μl of Edwards buffer is not easy, but still yields reliable results, while 5 ∼ 10 mg of leaf tissues is the optimal range for biomass input.

**Figure 2.**
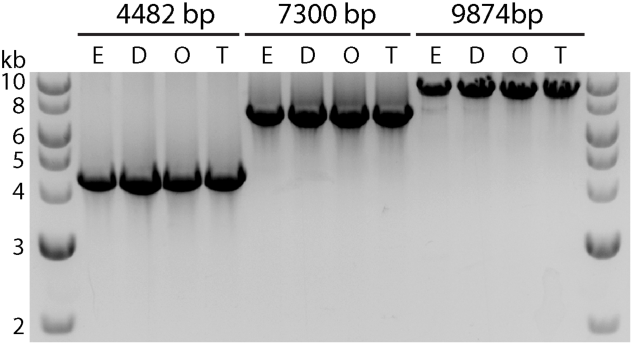
DNA extracted by the one-tube protocol is effectively amplified by long-range PCR using LongAmp Taq DNA polymerase. Two independent DNA extracts using Edwards (ED) and one-tube (OT) protocols are shown.

**Figure 3.**
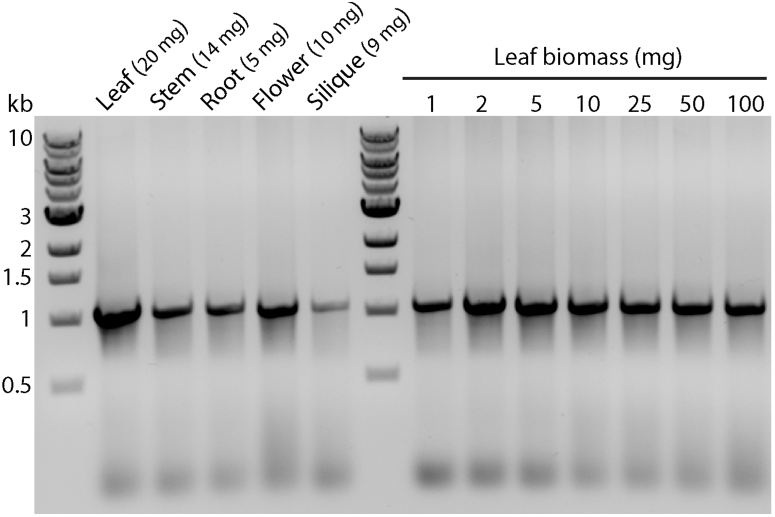
DNA extracts using the one-tube protocol from various Arabidopsis tissues and leaf biomass have negligible inhibitory effect on PCR analysis. For leaf biomass and size comparison, 1-2 mg is approximately equivalent to one cotyledon; 5 mg is equivalent to the leaf mass excised using the inner cap of a 1.5 ml microcentrifuge tube; 25-50 mg is equivalent to one single fully developed rosette leaf.

We have employed this one-tube method for routine PCR genotyping of *A. thaliana* plants. This method was also successfully used for isolation and PCR analysis of genomic DNA from transgenic tobacco and tomato plants (data not shown).

## Conclusions

The one-tube DNA extraction method described here significantly reduces extraction time, while maintaining the same DNA quality (as judged by PCR analysis) compared to that of the original Edwards method [1]. Moreover, despite the presence of cellular debris, one-tube DNA extracts are stable at room temperature for at least three months. This method is simple, rapid, robust, inexpensive and environmentally friendly. Plant biologist can benefit from the reduction in both time and cost when using this DNA extraction method for PCR analysis.

## Materials and Methods

### DNA extraction

The one-tube plant DNA extraction protocol developed in this study is described below. For the one-step extraction protocol of Kasajima *et al.* [3], the ground tissue solution was further centrifuged briefly to precipitate debris, which we found to improve the success rate for PCR reactions.

#### One-tube DNA extraction protocol

1. Collect ∼5-15 mg of leaf tissue into a 1.5 ml microcentrifuge tube;
2. Add 200 μl Edwards buffer [1], grind well with a plastic pestle (< 20 sec);
3. Add 200 μl isopropanol and mix well by gentle inversion;
4. Centrifuge 13,000 rpm x 5 min;
5. Decant the supernatant, invert the tube on paper tower to air dry the pellet (< 2 min);
6. Add 100 μl of sterile deionized H2O, drag the tube through a tube rack to quickly suspend the pellet (more effective and easier than vortex);
7. Centrifuge 13000 rpm x 1 min to precipitate insoluble debris;
8. Take 1-1.5 μl of supernatant for 20 μl of PCR analysis.

DNA solutions extracted with the above method are stable for at least three months at room temperature and at least one year at 4°C for reliable PCR analysis.

### Primers used in this study

Primers for PCR analyses priming the *A. thaliana PHYB* gene were oWH13 (5’-CAACAGCGGGAACAATGAAATG-3’), oWH14 (5’-TGAACGCAAATAATCTCCCTCTTC-3’), oWH31 (5’-TGTTTAAGCAGAACCGTGTCCG-3’), oWH32 (5’-TTCCATCCATTGATGCAGCCTC-3’), oWH105 (5’-GCGATTGGTGGCCAAGAT-3’), oWH130 (5’-TGATTCACCCTAAATCCTTCCTTG-3’), and oWH131 (5’-TGCTGTGTGCGGTATGGCAG-3’). oWH131 and oWH32 were for amplifying a 438 bp fragment, oWH31 and oWH32 for 1099 bp fragment, oWH105 and oWH14 for 1542 bp fragment, oWH130 and oWH32 for 2109 bp fragment, and oWH130 and oWH13 for 3535 bp fragment, respectively. The primers used for long-range PCR were chosen for priming a genomic *PHYB*^*Y276H*^ transgene allele (*YHB*^*g*^*/phyA-201 phyB-5* Line #5, [4, 5]); they were oWH130 and oWH197 (5’-TGCTCTAGCATTCGCCATTCAG-3’) for amplifying 4482 bp fragment, oWH130 and oWH40 (5’-CACCGACTACGCTTCACAGAAAG-3’) for 7300 bp fragment, and oWH120 (5’-TGAAGGCGGGAAACGACAATC-3’) and oWH40 for 9874 bp fragment.

### PCR amplification and gel electrophoresis

For routine PCR, 1 ∼ 1.5 μl of DNA extract was added to 20 μl of PCR reaction mix containing standard PCR buffer and 1 unit of *Taq* DNA polymerase (Cat # = M0273, New England Biolabs, Beverly, MA). For long-range PCR, 1.5 μl of DNA extract was added to 25 μl of LongAmp Taq DNA polymerase PCR reaction mix (Cat # = M0323, New England Biolabs) following the manufacturer’s protocol. PCR reactions were performed using a DNA Engine thermal cycler (Bio-Rad, Hercules, CA) with optimized annealing temperature for 35 ∼ 40 cycles. All 20 μl of routine PCR products and 2.5 μl of long-range PCR products were loaded for 1% agarose gel electrophoresis. Gel images were inverted in Adobe Photoshop.

## Authors’ contributions

W.H. designed and performed all experiments; J.C.L. provided funding, facilities and supervision. Both authors have read, corrected and approved the final version of the manuscript.

## Acknowledgements

We thank Timothy Butterfield for critical reading of the manuscript. This work was supported by grant GM068552 from the National Institutes of Health.

